# *Zymobacter palmae* pyruvate decarboxylase is less efficient than that of *Zymomonas mobilis* for ethanol production in metabolically engineered *Synechocystis* sp PCC6803

**DOI:** 10.1101/701193

**Authors:** Lorraine Quinn, Patricia Armshaw, Tewfik Soulimane, Con Sheehan, Michael P Ryan, J Tony Pembroke

## Abstract

Pyruvate decarboxylase (PDC) from *Zymobacter palmae* (ZpPDC) has been reported to have a lower Km the *Zymomonas mobilis* PDC (ZmPDC). ZpPDC was combined with native *slr1192* alcohol dehydrogenase (*adh*) in an attempt to increase ethanol production in the photoautotrophic cyanobacterium *Synechocystis* sp. PCC 6803 over constructs created with Zm*pdc*. Native (Zp*pdc*) and codon optimised (ZpO*pdc*) versions of the ZpPDC were cloned into a construct where the *pdc* expression was controlled via the *psbA2* light inducible promoter from *Synechocystis* PCC 6803. These constructs were transformed into wildtype *Synechocystis* PCC 6803. Ethanol levels were then compared with identical constructs containing the Zm*pdc*. While strains with the Zp*pdc* (UL071) and ZpO*pdc* (UL072) constructs did produce ethanol, levels were lower compared to a control strain (UL004) expressing the *pdc* from *Zymomonas mobilis*. The utilisation of a PDC with a lower Km from *Zymobacter palmae* did not result in enhanced ethanol production in *Synechocystis* PCC 6803.

## Introduction

Recently, much effort has focused on the development of alternative sources of energy that are environmentally friendly and sustainable [1] in comparison to fossil fuels [2, 3]. With many alternatives being explored, much research has focused on the metabolic engineering of the model cyanobacterium *Synechocystis* sp. PCC 6803 to produce biofuels such as ethanol [4] from CO_2_ and sunlight in a similar manner to plants [5]. Metabolic engineering has been utilised to direct *Synechocystis* PCC 6803 to produce a range of products [6] including ethanol via expression of heterologous *pdc* (pyruvate decarboxylase) and *adhII* (alcohol dehydrogenase) genes from *Zymomonas mobilis* (Zm*pdc* and Zm*adhII*).

In the late 1980s, *Escherichia coli* was initially engineered with these genes [7] to produce ethanol as a proof of concept. This was followed by the metabolic engineering of the first cyanobacterium *Synechococcus elongatus* sp. PCC 7942 to produce ethanol [8] and soon afterwards *Synechocystis* PCC 6803 was also used to produce ethanol using the same *pdc* and *adhII* genes from *Zymomonas mobilis* but using a strong light driven native *psbA2* promoter. This strain gave double the amount of ethanol production compared to the *Synechococcus elongatus* sp. PCC 7942 strain [9]. US biofuel companies Algenol and Joule Unlimited have since worked towards the development of industrial ethanol producing cyanobacteria using an *adh* (slr1192) native to *Synechocystis* PCC 6803 coupled to the *pdc* gene from *Zymomonas mobilis* allowing overexpression of these genes and enhanced ethanol production [10]. To enhance ethanol production further, gene dosage has been used which employed two copies of the Zm*pdc* and slr1192 *adh* genes coupled with the knockout of the PHB (poly-β-hydroxybutyrate) storage compound pathway [11] leading to further increased ethanol yields. Other approaches which hold potential include the use of small native *Synechocystis* PCC 6803 plasmids for expression of the heterologous genes in the ethanol pathway [12], the alteration of pyruvate levels via the over expression or decreased expression of certain enzymes like pyruvate kinase (PK) or phosphoenolpyruvate carboxylase (PPC) [13,14] or the utilisation of different promoters [15] for expression of the *pdc* and *adh* genes [14] to enhance enzyme activity and flux to ethanol.

We hypothesised that the *Zymomonas mobilis pdc* gene could be replaced with a *pdc* gene from *Zymobacter palmae* that possesses a PDC with a reported lower Km value. This could potentially increase flux from pyruvate to ethanol in engineered *Synechocystis* PCC 6803 strains. This possibility was examined via cloning and expression of the Zp*pdc* gene, both native (Zp*pdc*) and codon optimised (Zp*Opdc*), with these cassettes compared to those expressing the Zm*pdc* with respect to ethanol.

## Methods

### Bacterial Strains

The bacterial strains, plasmids and DNA elements utilised as part of this study are listed in Table 1. *Synechocystis* PCC6803 (glucose tolerant, obtained from K. Hellingwerf, UvA, Amsterdam) cells were maintained at 30°C on BG-11 media (Sigma) supplemented with 10 mM TES-NaOH (pH 8.2), 20 mM glucose and 0.3 % (w/v) sodium thiosulfate. *Zymobacter palmae* DSM-10491 was obtained from the DSMZ and grown in MY broth (1 g yeast extract, 2 g maltose [20% solution made, filter sterilised, 10 ml added for 2% after autoclaving], 0.2 g kH2PO4, 0.5 g NaCl). All routine plasmid construction and cloning was performed in *E. coli* using Luria–Bertani (LB) broth. All media were supplemented with appropriate antimicrobial agents as required: ampicillin, 100 µg ml^−1^ and kanamycin, 5–100 µg ml^−1^. All strains were stored at -80C in either Luria–Bertani (LB) broth containing 50 % glycerol (*E. coli*) or 50 % BG-11 media containing 5 % (v/v) methanol (*Synechocystis* PCC6803).

**Table 1:** Bacterial strains and plasmids used in this study

| Strain | Genotype/phenotype | Source |
| --- | --- | --- |
| BL21 (DE3)* | *E. coli* B F<sup>-</sup> *ompT*, *gal*, *dcm*, *lon*, *hsdS<sub>B</sub>*(*r<sub>B</sub><sup>-</sup>m<sub>B</sub><sup>-</sup>*), $\lambda$(DE3 [*lacI lacUV5-T7p07 ind1 sam7 nin5*]) [*malB<sup>+</sup>*]<sub>K-12</sub>($\lambda^S$) | Thermo Fisher Scientific Ballycoolin, Dublin 15, Ireland |
| DSM-10491 | *Zymobacter palmae* | DSMZ - German Collection of Microorganisms and Cell Cultures GmbH |
| AA314 | Wild-type *Synechocystis* PCC6803 strain | K. Hellingwerf, UvA, Amsterdam, The Netherlands |
| UL004 | PCC6803 transconjugant with ZmPDC, *slr1192- adh*, *psbA2* locus, Kan<sup>R</sup> | [19], This study |
| UL070 | PCC6803 transconjugant with ZmPDC, *slr1192- adh*, *psbA2* locus, Kan<sup>R</sup> | This study |
| UL071 | PCC6803 transconjugant with ZpPDC, *slr1192- adh*, *psbA2* locus, Kan<sup>R</sup> | This study |
| UL072 | PCC6803 transconjugant with ZpOpdc, *slr1192- adh*, *psbA2* locus, Kan<sup>R</sup> | This study |
| Plasmid | Genotype/phenotype | Source |
| pUC18 | Amp<sup>R</sup> backbone plasmid | Sigma-Aldrich, Arklow, Wicklow, Ireland |
| pUL004 | pUC18 backbone, *PpsbA2* promoter, *ZmPDC*, *slr1192- adh*, *psbA2* integration site, Kan<sup>R</sup> | [19] |
| pUL101 | pUC18 backbone *PpsbA2* promoter, *ZpPDC*, *slr1192- adh*, *psbA2* integration site, Kan<sup>R</sup> | This study |
| pUL102 | pUC18 backbone *PpsbA2* promoter, *ZpOpdc*, *slr1192- adh*, *psbA2* integration site, Kan<sup>R</sup> | This study |

### Gene Cloning and Strain Construction

Cloning via homologous recombination was carried out using the In-Fusion® HD cloning kit (Clontech Laboratories Inc). Primers used can be seen in Table 2.

**Table 2:**
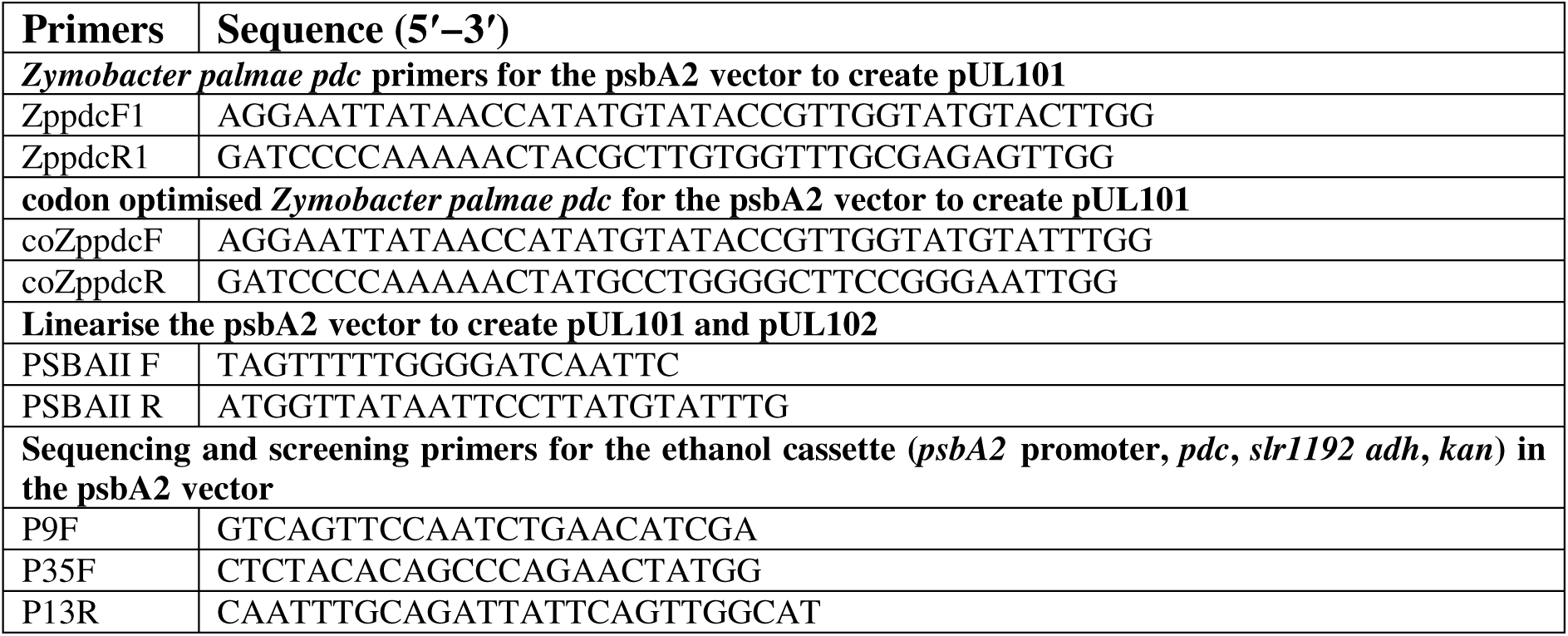
Primers used in this study

Transformation was carried out via electroporation with electro-competent cells [16]. The gene sequence for *Zppdc* was sent to IDT (Integrated DNA Technologies) to be codon optimised for *Synechocystis* 6803. Cassettes were constructed which contained the pPSBAII light promoter (from plasmid pUL004, Table 1) fused to the Zm*pdc*, the Zp*pdc* and the ZpO*pdc* genes coupled to the native *Synechocystis* PCC 6803 slr1192 *adh* gene and the *kanamycin* resistance gene derived from the ICE R391 [17] as per Lopez et al [18] (Fig. 1). The construct also contains 500 bp at each end with homology to the PSBAII neutral site to allow homologous recombination into the PSBAII neutral site [10]. Constructs were generated by PCR amplification of the relevant genes and promoter with fusion of the genes carried out via a biobrick to form the ethanol cassettes similar in structure as previously reported [9]. Verification of the pUL101 and pUL102 plasmids was carried out via PCR amplification of the construct (Table 2) followed by sequencing. PCR mixes, Taq Polymerase and restriction enzymes for cloning and PCR methods were purchased from Sigma Aldrich. PCR cycle was as follows: after an initial denaturation at 98 °C for 1 min, 30 cycles of denaturing at 98 °C for 10 s, annealing at 50 °C for 20 s and extension at 72 °C for 1 min (30 secs/kb for ∼2kb gene) extension were undertaken, with a final step at 72 °C for 5 min. PCR products were then analysed by electrophoresis on a 1.0 % agarose gel stained with Sybersafe using 1X TAE buffer as running buffer.

**Fig 1:**
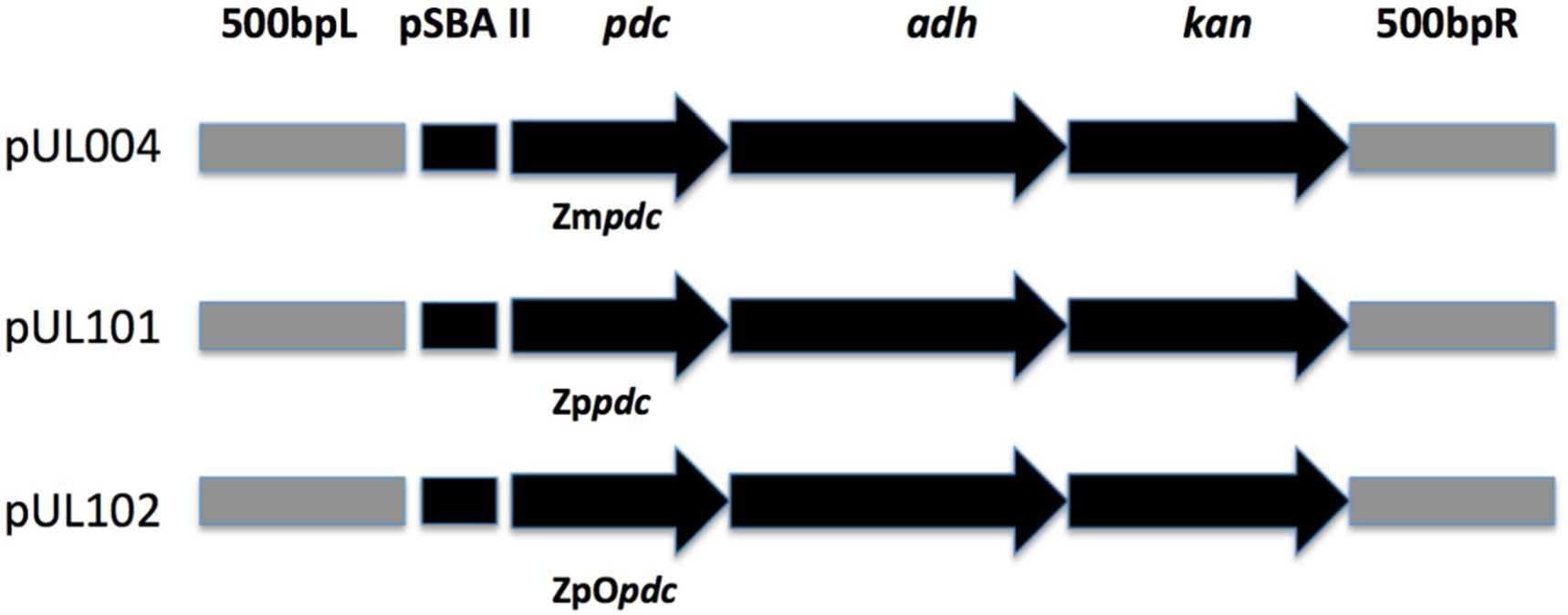
Structure of the recombinant cassettes transformed in *Synechocystis* sp. PCC 6803.

Transformants of wild type *Synechocystis* were sub-cultured in BG-11 media containing increasing concentrations of kanamycin [5–100 µg.ml^−1^] until full integration of the cassette was verified. Transformations were left at 30 °C for 16 h under medium intensity white-light illumination (∼20–40 µE m^−2^ s^−1^). Verification of integration into the *psbA2* neutral site was carried out with appropriate primers (Table 2) that bound the flanking homologous insertion site within psbA2. Wildtype *Synechocystis* amplified with these primers generated a PCR product approximately 1.2 kb in size, insertion of the UL071 and UL072 cassette resulted in the amplification of a ∼4 kb PCR product.

### Growth Measurements

Optical density measurements were taken using a Cary UV-Vis spectrophotometer at either 600nm for *E.coli* or 730nm for *Synechocystis* sp. PCC 6803.

### Ethanol Determination

Ethanol determination was carried out using the Yellow line kit: UV-method from R-Biopharm AG. Here, ethanol is oxidised to acetaldehyde via alcohol dehydrogenase (ADH) which in the presence of aldehyde dehydrogenase (Al-DH) is oxidised to acetic acid while NAD^+^ is reduced to NADH, which was measured at 340 nm via a Cary UV-Vis spectrophotometer. All tests were carried in quintuplicate.

## Results and Discussion

Bacterial PDCs are rare with only a small number reported. Raj *et al.* reported four bacterial PDCs from *Zymomonas mobilis* (Zm), *Zymobacter palmae* (Zp), *Acetobacter pasteurianus* (Ap) and *Sarcina ventriculi* (Sv) [20]. By aligning the amino acid sequences of these PDCs which are similar in length and size (between 552 and 568 amino acids and 59.83 and 61.8 KDa) it was found that the PDC from *Z. palmae* (ZpPDC) shared 72% identity with the PDC from *Acetobacter pasteurianus* (ApPDC) but only 62/63% identity to the ZmPDC [Table 3, 20, 21]. Comparison of kinetic parameters, which can be seen in Table 3, indicated that the ZpPDC might have some potential in metabolic engineering due to its lower Km value in comparison to the ZmPDC. Two other PDC’s have since been reported from *Gluconaacetobacter diazotrophicus*, GdPDC, and *Gluconobacter oxydans*, GoPDC [22] and are compared in Table 3.

**Table 3:** Characteristics of known bacterial Pyruvate Decarboxylases

| PDC | ZmPDC | ZpPDC | ApPDC | SvPDC | GoPDC | GdPDC |
| --- | --- | --- | --- | --- | --- | --- |
| **PDB entry** | 1ZPD | 5EUJ | 2VBI | N/A | N/A | 4COK |
| **Bacterial species** | *Zymomonas mobilis* | *Zymobacter palmae* | *Acetobacter pasteurianus* | *Sarcina ventriculi* | *Gluconobacter oxydans* | *Gluconacetobacter diazotrophicus* |
| **Gram** | Negative | Negative | Negative | Positive | Negative | Negative |
| **Gene** | M15393 | AF474145 | AF368435 | AAL18557 | KF650839 | KJ746104 |
| **Protein** | AAA27696 | AAM49566 | AAM21208 | AF354297 | AHB37781 | AIG13066 |
| **Amino acid identity %** | *62/^63 | Reference | ^73 | ^31 | ^67 | ^71 |
| **Kinetics** | *M-M | *M-M | *M-M | * Sigmoidal | *M-M | *M-M |
| **Km mM (pH)** | *0.43(6.0)<br>0.94 (7.0) | *0.24 (6.0)<br>0.71 (7.0) | *0.39 (5.0)<br>5.10 (7.0) | *5.7 (6.5)<br>4.0 (7.0) | #0.12 (5.0)<br>1.20 (6.5)<br>2.80 (7.0) | #0.06 (5.0)<br>0.60 (6.0)<br>1.20 (7.0) |
| **Optimum Temperature °C** | *60 | *55 | *65 | N/A | #53 | #45-50 |
| **Optimum pH** | *6.0 | *5.5-6.0 | *5.0-5.5 | *6.3-6.7 | #4.5-5.0 | #5.0-5.5 |
| **Reference** | [25] | [21] | [26] | [20] | [20] | [22] |

M-M: Michaelis-Menten

pUL004 (containing Zm*pdc*), pUL101 (Zp*pdc*) and pUL102 (ZpO*pdc*) (Table 1) were transformed into wildtype *Synechocystis* PCC 6803 to create strains UL070 (pUL004), UL071 (pUL101) and UL072 (pUL102) respectively. The Zppdc gene sequence was codon optimised to minimise the effect of codon bias in limiting expression in the heterologous host giving UL072. Full segregation of the constructs into the chromosome at the psbA2 neutral site was confirmed via PCR screening. Using primers that spanned the *psbA2* insertion site, wildtype strains or those that failed to integrate cassettes into the polyploidy chromosome resulted in an amplicon of ∼1.1 kb. Fully segregated strains that integrated the cassettes also showed one band but at a size of ∼4 kb (containing the *psbA2* light promoter, *pdc, adh* and *kanamycin* genes). Strains that displayed partial integration into only some of the polyploid chromosomes showed two bands of both 1.1 and 4 kb (Supplementary Material).

It was decided to test overall ethanol levels as increasing levels of ethanol was the desired outcome of the research. All strains produced 0 g/L/OD of ethanol on day 0 and levels for each strain subsequently varied over the course of the 3, 7 and 11 days upon growth in BG11 media. UL070 containing Zm*pdc* produced the largest amount of ethanol at each measurement time in comparison to the other strains (Fig 2).

**Fig 2:**
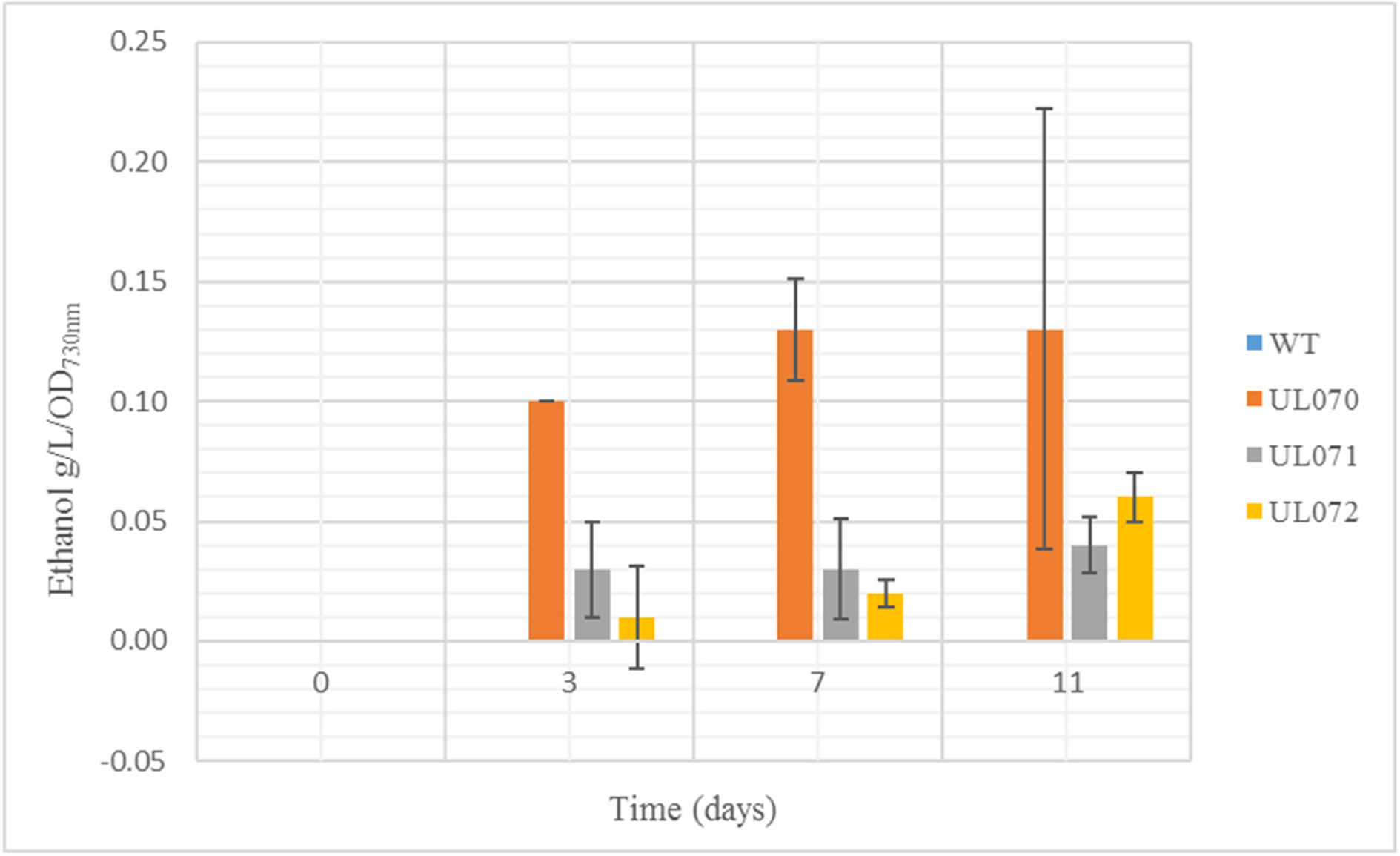
Ethanol levels in g/L/OD730nm for tested WT, UL070, UL071 and UL072 strains on days 0, 3, 7 and 11 (*n*=5)

Ethanol levels for the UL071 and UL072 strains were less than the Zm*pdc* expressing strain which was somewhat surprising, given the lower Km that has been reported for this PDC (we subsequently purified the ZpPDC and verified its reported lower Km, data not shown) [20]. All recombinant strains grew at a slower rate than wildtype that is typical of strains diverting key metabolic intermediates such as pyruvate away from biomass and other metabolic needs. This can be observed in differential Optical Density (OD) in ethanol producers relative to wildtype strains. All three strains examined showed reduced OD after 3, 7 and 11 days of culture relative to the wildtype *Synechocystis* PCC 6803 strain with UL070 showing the lowest OD which is an indication that it was most effected by the ethanol production relative to biomass (Fig 3).

**Fig 3:**
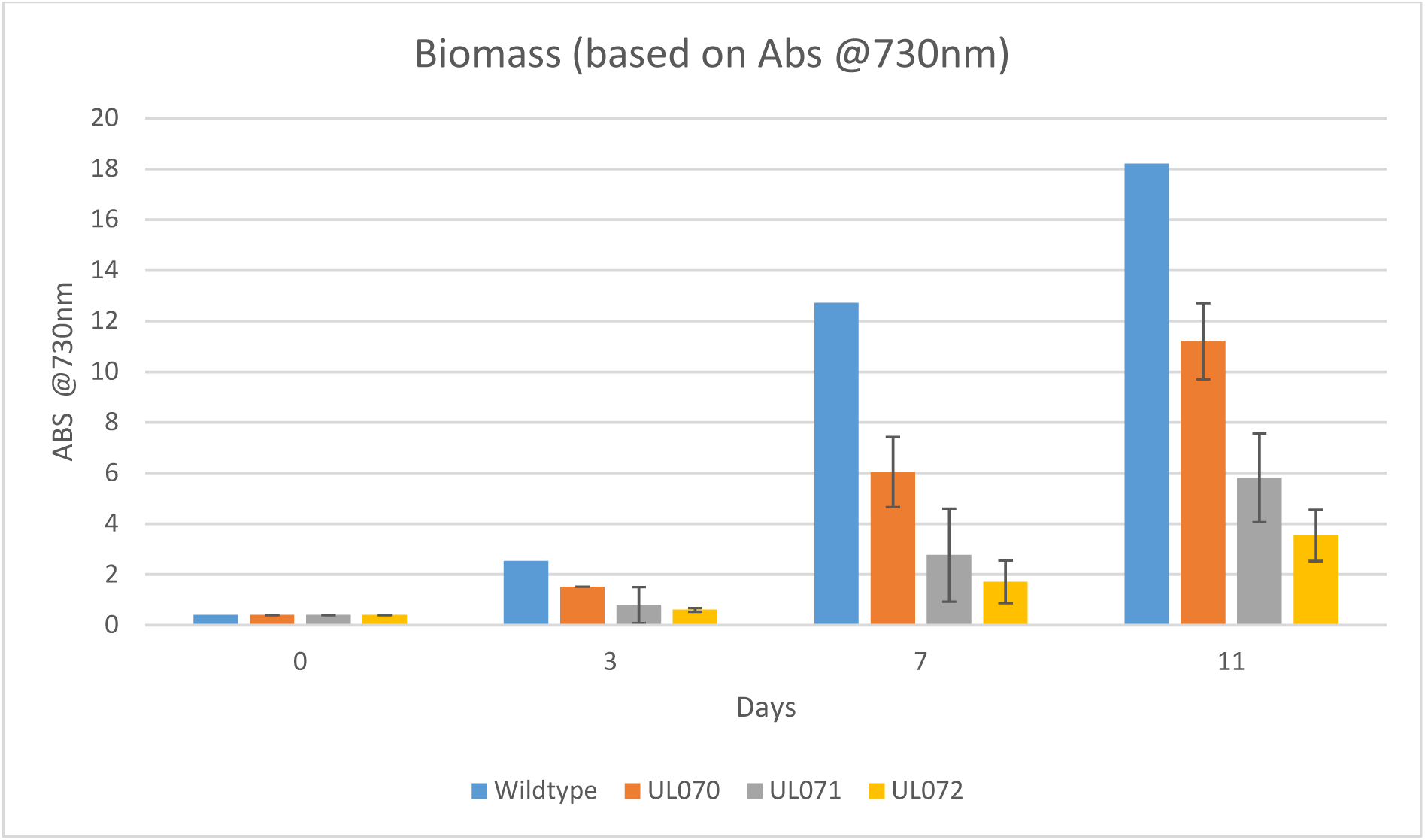
Biomass levels (OD730nm) for tested WT, UL070, UL071 and UL072 strains on days 0, 3, 7 and 11 (*n*=5)

Liu *et al*. also used Zp*pdc* in a study to increase ethanol production in lactic acid bacteria [23]. The Zp*pdc* was chosen for the same reasons as outlined above: its low Km for pyruvate and high specific activity amongst all the bacterial PDCs [24]. By using acid inducible and highly conserved constitutive promoters with the Zp*pdc*, Liu *et al*. reported that the acetaldehyde levels produced by the recombinant strains of *Lactococcus lactis* were eight fold higher in comparison to the control strain but that there was no significant increase in ethanol levels [23]. There may be several reasons why the Zp*pdc* is not performing as might be hypothesised. It is possible that Zp*pdc* is converting pyruvate at a faster rate to acetaldehyde compared to the Zm*pdc* and that this is not coupled effectively to the ADH resulting in build-up of acetaldehyde that is known for its toxicity [24]. It may also be possible that there is a pH incompatibility issue as the optimum pH for ZpPDC is pH 6.0 [20] which is slightly more acidic than the optimum pH for *Synechocystis* PCC 6803 which is 8.2 with growth showing little reduction up to pH10 (26) (At this pH ZpPDC displays ∼60% of the activity displayed at pH 6.0). In addition to this, the ZpPDC has received little study and so it is possible that there are cofactor differences, metabolic regulatory issues or coupling issues, which have yet to be recognised. Although utilising a PDC with a lower Km has potential, our data indicates that before such potential can be realised more detailed studies on candidate PDCs will be necessary before progress in this area can be achieved. The utilisation of a PDC with a lower Km from *Zymobacter palmae* in both the native and codon optimised form in a pathway from ethanol formation in *Synechocystis* PCC 6803 did not result in an increase of ethanol production levels under the conditions tested.

## Author Contributions

J.T.P, P.A, L.Q., T.S C.S. conceptualized, planned, designed, analysed the study and interpreted results M.R. interpreted results and wrote the manuscript.

