## Supplementary Figure and Table for "*Zymobacter palmae* pyruvate decarboxylase is less efficient than that of *Zymomonas mobilis* for ethanol production in metabolically engineered *Synechocystis* sp PCC6803"

### Supplementary Material

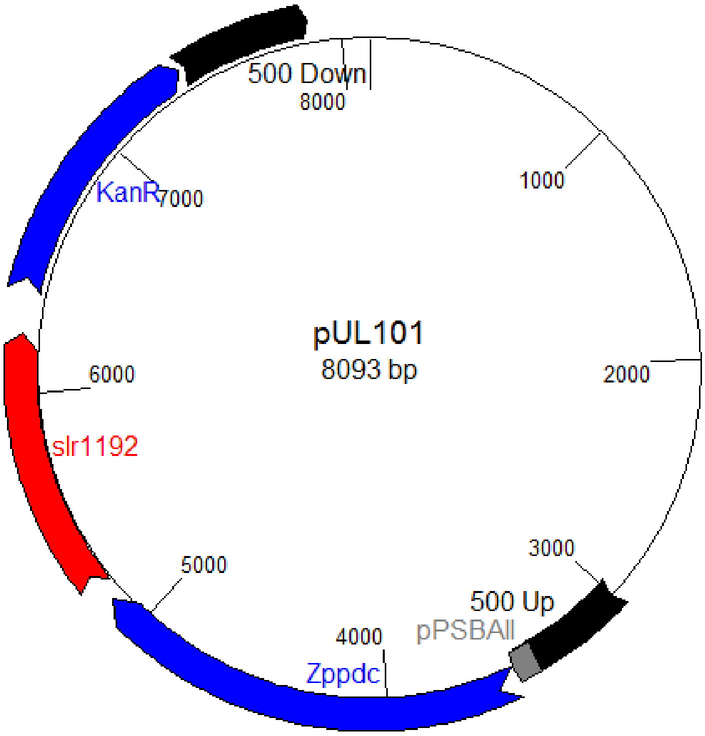


**Figure 1: pUL101** This construct shows the *psbA2* promoter of *Synechocystis* 6803 amplified and fused to the native *Zymobacter palmae pdc* (Zp*pdc*) and linked to the native *Synechocystis* 6803 *adh* (slr1192) and *kanamycin* resistance gene (a gene encoding kanamycin phosphotransferase which was amplified from the enterobacterial ICE R391). The construct that was verified by sequencing contained two 500 bp flanking regions that were homologous to the *psbA2* neutral site in *Synechocystis* 6803. It was now suitable for transformation and incorporation into this site. Transformants were initially screened for kanamycin resistance with the level of kanamycin gradually increased to allow for screening and selection for full segregation into the polyploidy genome of *Synechocystis* 6803. Primer sets spanning the *psbA2* site were designed to differentiate between full incorporation of this construct into every copy of the chromosome, partial non-segregated incorporation and no incorporation.

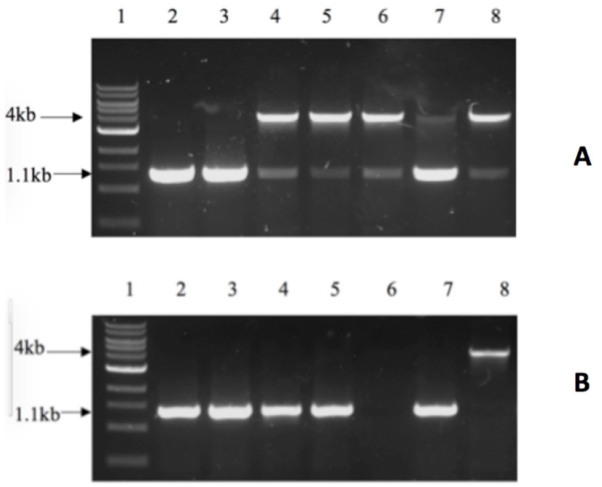


**Fig 2:** 1% DNA agarose gel of PCR amplicons. Molecular weight markers are shown in L1 for both A and B. A: wildtype (L2, 1.1 kb band), unsegregated UL072 strains (L3 1.1 kb band), semi-segregated UL072 strains (L4-L8, 1.1 and 4 kb bands). B: wildtype (L2, 1.1 kb band), unsegregated UL071 strains (L3-L5, 1.1 kb bands), empty (L6, no band), unsegregated UL071 strain (L7, 1.1 kb band), fully segregated UL071 strain (L8, 4 kb band only).

**Table 1:** Ethanol yields as measured in g/L/OD for wild-type and metabolically engineered strains (*n*=5).

| **Time (days)** | **Wildtype** | **UL070 (Zm*pdc*)** | **UL071 (Zp*pdc*)** | **UL072 (ZpO*pdc*)** |
|---|---|---|---|---|
| 0 | 0.00 | 0.00 | 0.00 | 0.00 |
| 3 | 0.00 | 0.10 ±0.00 | 0.03 ±0.02 | 0.01 ±0.02 |
| 7 | 0.00 | 0.13 ±0.02 | 0.03 ±0.02 | 0.02 ±0.01 |
| 11 | 0.00 | 0.13 ±0.09 | 0.04 ±0.01 | 0.06 ±0.01 |
